# Stomatal setpoints and environmental responsiveness are sculpted by developmental trajectories

**DOI:** 10.64898/2025.12.22.696041

**Authors:** Monalisha Rath, Nidhi Sharma, Madhav Mani, Dominique C. Bergmann

## Abstract

Efficient gas and water exchange between plants and their environment largely depends on the number and distribution of stomata, cellular valves in leaf epidermis. Core genetic regulators of stomatal cell identity and pattern along with asymmetric stem-cell like divisions in stomatal precursors are hypothesized to customize stomatal production for optimal leaf performance. How these regulators work in concert and how division dynamics are modified and adjusted in different environments, however, are poorly understood. Here, we leveraged the variation in stomatal patterning in Arabidopsis thaliana accessions from diverse environments to define developmental rules and constraints in the stomatal lineage. The accessions’ subtle and quantitative variation enables us to identify which cellular parameters are flexible, revealing how developmental plasticity generates phenotypic plasticity. By developing live-cell imaging tools to track cellular behaviors during leaf growth under varying environmental conditions in these accessions, we could decompose stomatal density variation into its developmental origins. Variation in final stomatal numbers is driven by differences in the relative contributions of stomatal initiation, cell size-based fate thresholds, general proliferative capacity, and coordination between sister and neighbor cell behaviors. Overall, diverse accessions converge toward two lineage regimes: one dominated by autonomous decisions with loose cell-cell coordination, the other by extensive cell-cell coordination. Challenging accessions with environmental fluctuations revealed regime-specific flexibility, with plasticity primarily mediated by a single division-related parameter. Our results show how cellular parameters integrate into alternative developmental strategies that shape environmental responsiveness.

## INTRODUCTION

Phenotypic plasticity is the ability of a genotype to yield different morphological, physiological, or behavioral outcomes when exposed to distinct environmental conditions. The self-fertile and highly inbred dicot plant *Arabidopsis thaliana* has served as a powerful model to explore phenotypic plasticity, its evolutionary drivers and its ecological consequences (reviewed in ^1–3^). Because plants can create new organs throughout their lifetime, plasticity often manifests in the number, size, shape and cellular composition of these organs. These organ traits are typically measured at maturity ^4^, but endpoint measurements can obscure variation in the developmental pathways leading there. This limitation is particularly acute because a single mature phenotype can arise through distinct developmental routes that differ in cost, robustness and adaptability of the plant ^5^. For example, smaller leaves produced in response to drought stress could result from reduced cell proliferation, premature cessation of cell division, decreased cell expansion, increased cell death, or any combination of these processes ^6,7^. Moreover, endpoint analyses may entirely miss compensatory adjustments, where multiple parameters change in opposing directions such that their effects on the final phenotype cancel out ^8^. The cellular dynamics underlying plasticity, including which specific decision points are environmentally sensitive, which are buffered, and how they are coordinated has been explored in the highly stereotyped development of seedling hypocotyls and roots ^9,10^. The extent to which the mechanisms uncovered in these studies apply to tissues where multiple developmental paths can be taken to reach a common endpoint, remains largely unknown.

It has been long observed that leaves change in size and shape and in the production and distribution of specialized vascular and epidermal cells in response to the environment ^11,12^. These phenotypic changes emerge when plants are shifted to new locations, or even when comparing leaves from sun-exposed and shaded areas of the same tree ^11^. The stomatal lineage in the leaf epidermis offers a tractable system to correlate phenotypic outcomes with the cellular and developmental steps used to get there. Dicot stomatal lineages exhibit substantial developmental diversity^13–15^, and studies in *Arabidopsis* provide a well-characterized framework in which a series of stereotyped yet flexible cell division patterns and fate acquisition events precede stomatal differentiation. Dispersed stem-cell like precursor cells initiate stomatal lineages through an asymmetric “entry” division, creating meristemoids (M) as their smaller daughters and stomatal lineage ground cells (SLGCs) as their larger daughters (Fig. 1A) ^16^. Both meristemoids and SLGCs can undergo additional rounds of asymmetric cell divisions, termed amplifying and spacing divisions, respectively. Each division produces more cells in the epidermis: amplifying divisions decrease the ratio of stomata to non-stomatal cells whereas spacing divisions increase stomatal production relative to non-stomatal cells. In both Ms and SLGCs, cell size serves as a differentiation cue, and cells that reach a threshold size differentiate ^17,18^. Flexibility in division behaviors has been proposed to enable leaves to optimize gas exchange, transpiration and photosynthesis to a wide range of environmental conditions ^19,20^. Because the divisions in the stomatal lineage produce the majority of epidermal cells ^21^ and the epidermis guides overall organ growth ^22^, regulating stomatal lineage progression has a major overall effect on leaf development.

**Figure 1.**
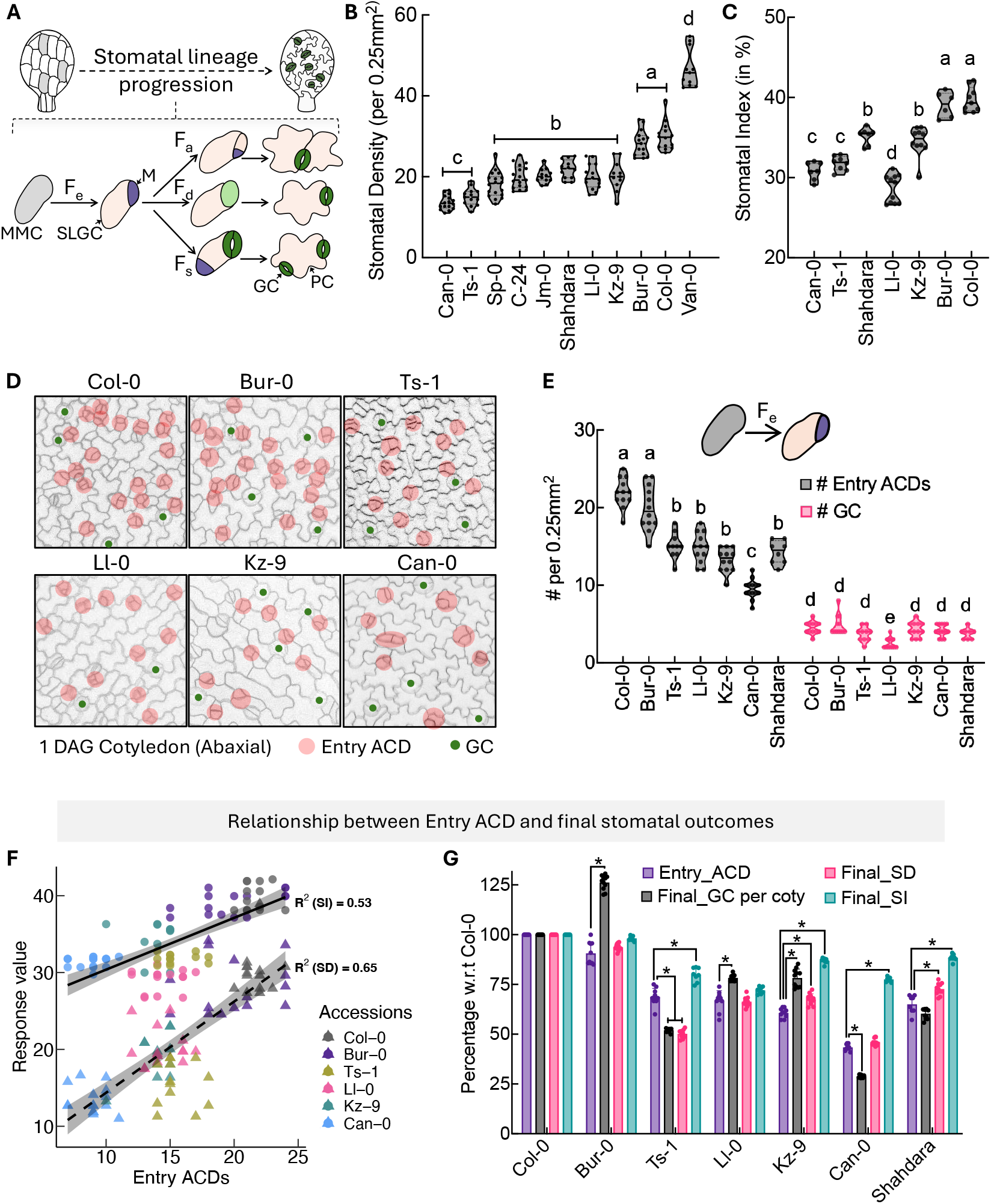
Natural accessions vary in stomatal lineage progression and overall stomatal production. A. Cartoon of asymmetric division types during stomatal lineage progression showing the lineage entry event (F_e_) of an meristemoid mother cell (MMC, grey), self-renewing amplification division (F_a_) or differentiation (F_d_) of the meristemoid (M, purple) and the spacing division (F_s_) of the SLGC producing mature guard cells (GC, green) and pavement cells (PC). An expanded version of the stomatal lineage with additional parameters measured in this paper is shown in Fig. 5C. **B-C**. Violin plots showing Stomatal Density (Final_SD, stomata/area) **(B)** and Stomatal Index (Final_SI, ratio of GCs to total epidermal cells) **(C)** in mature (20 day after germination, DAG) abaxial cotyledons of 11 Arabidopsis accessions. Shape of the violin indicates the data distribution density and different letters indicate significant difference (p < 0.01). **D**. Micrographs of abaxial epidermis of 6 representative accessions; entry ACDs marked with pink circles and GCs with green dots in 1DAG cotyledons **E**. Quantification of **D** for entry ACDs (black) and already formed guard cells (pink). **F**. Simple linear regression with all accessions pooled together showing the relationship between entry events and SD (dotted line) or SI (solid line) with the R^2^ values written inside the plot. Dots (for SD) and triangles (for SI) are color-coded by accession with each point representing a replicate. **G**. Relationship between entry divisions (Entry_ACD) and stomatal production was measured three ways: (Final_SI), (Final_SD) and total number of GCs per cotyledon area (Final_GC per coty) and calculated with respect to Col-0 values and plotted together. Different relationships between entry divisions and stomatal production among accessions indicate divergence in cellular pathways between these endpoints. Ratios calculated from data in Fig. 1B and 1E. Details on sample sizes and statistical tests are provided in Table S2.

Classical genetic screens identified key stomatal development genes and enabled elucidation of the core transcriptional and signaling networks that produce functional, well-distributed stomata (reviewed in ^23^). Analysis of loss-of function mutations in major regulators, however, misses the quantitative variation in stomatal numbers that may be adaptive. In many plant species^24–28^, natural accessions harbor integrated variation across multiple loci, regulatory networks, and cellular parameters related to stomata. In Arabidopsis, there is extensive natural variation in stomatal density, size and distribution, including subtle yet biologically meaningful differences in environmental responsiveness ^29,30^. From this starting point, by tracking cellular dynamics across accessions and environments, it is possible to measure the landscape of stomatal development parameters and show how they translate into phenotypic plasticity.

In this study, we probe natural variation not just at phenotypic endpoints, but along entire developmental trajectories in developing leaves. We created fluorescent live-cell imaging reporters in six Arabidopsis accessions, and through quantitative measurements of stomatal lineage behaviors and tracking multiple lineages over time, we reveal how cellular parameters govern developmental plasticity. All accessions employed size-based fate decisions and the same core developmental strategy of using a combination of asymmetric entry, amplifying, and spacing divisions to generate and pattern stomata. Accessions differed, however, in their absolute cell size thresholds, proliferative capacity, relative emphasis on each division type, and degree of cell-cell coordination. Variation converged on a limited number of “developmental regimes” with distinct environmental responsiveness, where plasticity emerged from a specific subset of the features defining the regimes. Within regimes, division frequency and behaviors provides developmental flexibility, while the coordinated behavior of sister and neighboring cells imposes constraints limiting stomatal production. Taken together, our study revealed the levers and constraints within developmental trajectories that sculpt environmental responses, informing both targets and limitations for engineering climate-adaptive stomatal development.

## RESULTS

### Arabidopsis accessions display variation in stomatal development that is not captured by mature tissue measurements

To identify Arabidopsis accessions occupying a wide range of stomatal lineage behaviors, we selected accessions collected from diverse regions and climates (Fig. S1A, Table. S1) that had been previously described as exhibiting morphological variation in leaves (arapheno.1001genomes.org) (Fig. S1B-D). A few of these accessions had been further characterized at the cellular level, and display differences in cell size and stomatal number ^30^. In our standard growth conditions (see methods) these accessions exhibit wide variation in stomatal density (SD, stomata per area) on both adaxial and abaxial surfaces of the mature cotyledons, independent of their mature, fully expanded cotyledon size (Fig. 1B, S1E-F). Compared with survey of 330 accessions ^29^our chosen accessions sampled the range of SD seen in Arabidopsis. Our selected accessions Stomatal Index (SI), the ratio of stomata to all epidermal cells, a measure that more directly reflects the history of stomatal lineage decisions than SD, also varies (Fig. 1C), and across these accessions SD and SI are positively correlated (Fig. 1B-C, S1G). The positive correlation between SI and SD indicates the variation in stomatal abundance among accessions arises from developmental lineage decisions (more so than differential expansion of epidermal cells) prompting us to investigate the cellular dynamics and lineage progression underlying these final outcomes.

### Lineage tracing shows diverse utilization of ACD subtypes among accessions

Final phenotypic diversity emerges from variations in the preceding cellular behaviors during development. Previous work in Col-0 suggested that the number of cells entering stomatal lineages and/or the subsequent behavior of those cells undergoing further amplifying or spacing divisions would be major sources of variation (Fig. 1A) ^17,21^. To capture stomatal lineage trajectories by timelapse microscopy during development in multiple accessions, we generated transgenic lines of six accessions (Col-0, Bur-0, Ll-0, Kz-9, Ts-1 and Can-0) each expressing a fluorescent plasma membrane reporter under the control of an epidermis-specific promoter (*AtML1p:RCI2A-mCherry*). These six accessions were chosen to represent a broad range of stomatal densities (Fig. 1B), and span diverse climatic regions based on their native collection sites (Fig. S1H and Table S1). When compared to the 1,135 sequence accessions from the 1001 genomes collection, the six accessions represent a broad genetic diversity (Fig. S1I).

Using lines expressing the epidermal plasma membrane marker, we captured images of the entire abaxial cotyledon epidermis. Although few stomata are present at 1 day after germination (DAG), this stage is marked by a surge in new lineage initiation, as evidenced by asymmetrically dividing sister cell pairs. These earliest measurable divisions we refer to as “entry ACDs” (Fig 1D). Compared to Col-0, accessions that displayed a reduction in mature stomatal numbers showed a significant decrease in entry ACDs (Fig. 1D-E). To assess whether the frequency of entry ACDs alone could predict final stomatal outcomes, we quantified the relationship between entry ACDs and the resulting stomatal traits—SD and SI. Simple linear regression analysis revealed that entry ACDs account for approximately half to two-thirds of the observed variation in final stomatal numbers, whether using pooled individual measurements (R^2^ ~ 0.53-0.65; Fig. 1F) or genotype-level means derived from BLUE estimates (Best Linear Unbiased Estimators) (R^2^ = 0.64-0.85; Fig. S2A). Within-accession analyses showed even lower explanatory power (Fig. S2B-C). The residual plots confirmed model inadequacy, exhibiting systematic non-random patterns including a U-shaped trend and persistent accession-specific deviations (Fig. S2D-E), indicating that the relationship between entry ACDs and final outcomes is more complex than a simple proportional scaling. Comparative analysis across accessions revealed that some accessions (Kz-9, Shahdara) produced disproportionately high stomatal numbers relative to entry ACDs, while others with similar frequencies of entry ACDs (Ts-1, Ll-0) showed markedly different SD and SI values (Fig. 1G). Together, these results demonstrate that while entry ACDs contribute to stomatal variation, they alone are insufficient to explain the different developmental endpoints, prompting us to examine the involvement of other asymmetric division types and developmental controls.

Beyond the number of entry ACDs, a key determinant of final stomatal production is the meristemoid’s capacity to continue asymmetric divisions or undergo differentiation. A recent study established a critical birth size threshold below which the meristemoids cease self-renewing *amplifying* divisions (F_a_) and commit to terminal differentiation ^17^. We tested whether this size-thresholded differentiation phenomenon was present across accessions and whether variation in this mechanism could account for accessions’ differences in stomatal production. By tracking meristemoid size and behavior over 24-hour timecourses we found that all accessions exhibited size-dependent meristemoid fate determination (Fig. 2A). Three low-SD accessions (Kz-9, Ll-0, Can-0) exhibited larger meristemoid size differentiation thresholds compared to high-SD accessions (Col-0, Bur-0), with Ts-1 being a low-SD exception sharing Col-0’s threshold (Fig 2A). Previous work identified nuclear size as a more reliable predictor of division and fate behavior than cell size (Gong et al. 2023). Consistent with these predictions, we found that nuclear size paralleled cell size among the accessions tested (Fig 2B). These data confirm that size-dependent fate control operates in all accessions but can exhibit shifted thresholds. The functional consequences of these divergent size thresholds became apparent with the quantification of meristemoid division behavior. Since meristemoid size shrink with successive divisions ^31^, larger thresholds require fewer divisions to trigger differentiation. Accordingly, low-SD accessions with their higher size threshold showed more frequent meristemoid differentiation and fewer rounds of amplification divisions compared to Col-0 (Fig. 2C). Importantly, amplifying divisions generate sister SLGCs, which can either differentiate into pavement cells or undergo spacing divisions to produce new meristemoids. Thus, the differentiation size threshold of meristemoid influences final SD not only by limiting meristemoid self-renewal, but also by determining the size of the SLGC pool available for secondary stomatal production. Meristemoid size as the sole explanatory factor for stomatal production, however, is challenged by several observations. First, accessions with similar thresholds can produce divergent stomatal densities: Ts-1 and Col-0 share comparable size thresholds yet differ markedly in final stomatal counts (Fig. 2A and 1B). Second, fewer amplifying divisions should increase SI by producing fewer non-stomatal cells, yet Ll-0 and Kz-9 show lower indices than Col-0 despite having fewer amplifying divisions (Fig. 1C and 2C).

**Figure 2.**
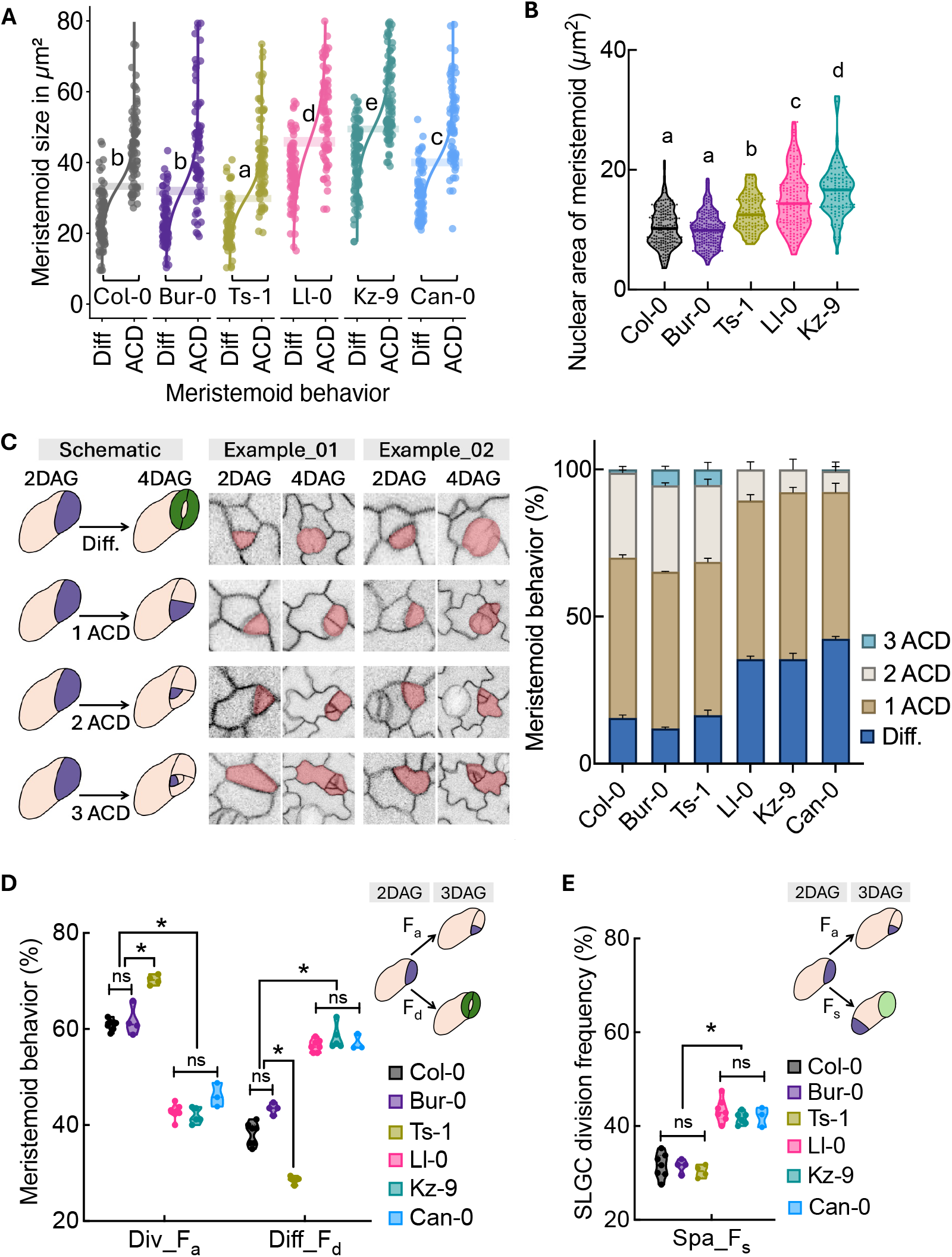
Meristemoid size impacts asymmetric divisions and phenotypic plasticity. A. Size distribution of meristemoids transitioning from division (ACD) to differentiation (Diff) at 2-3DAG. Logistic regression curves indicate the size-dependent differentiation of each accession, with the shaded region showing the transition size range where 50% of meristemoid differentiate. **B**. Nuclear area (as determined by Hoechst staining) of 3DAG meristemoids. Shape of violin indicates the data distribution density and different letters indicate significant difference (p < 0.01) **C**. (left) Examples of meristemoid division behaviors tracked from a time-course of 2-4 DAG cotyledons (right) quantification of division behaviors, n= 60 cells per cotyledon in each of 3 replicates. **D**. Frequency of dividing (Div., *f*_*a*_) or differentiating (Diff., *f*_*d*_) meristemoids in 2-3DAG cotyledons (24h time course). **E**. Frequency of SLGC division (*f*_*s*_) measured from the same cotyledons as in D. Details on sample sizes and statistical tests are provided in Table S2.

To resolve the above discrepancies, we examined the post-entry proliferative behaviors of both lineage branches: amplifying divisions in meristemoids and spacing divisions in SLGCs. By tracking the same leaves over a 24-hour period, we found that high-SD accessions (Col-0, Bur-0) showed significantly more amplifying than spacing divisions (60% vs. 30%), whereas low-SD accessions (Ll-0, Kz-9, Can-0) exhibited comparable frequencies of both division types (each ~40%, Fig. 2D-E). Since amplifying and spacing divisions have opposite effects on stomatal index, this divergence in division strategies explains why final stomatal numbers in Kz-9 and Can-0 are not as low as their entry division frequencies would predict (Fig. 1G). Correlating these behaviors with size thresholds revealed a noteworthy insight: accessions with lower meristemoid differentiation threshold size (high-SD accessions) showed ~20% more amplifying divisions, while those with higher thresholds (low-SD accession) favored spacing divisions (~10% more, Fig. 2D-E). Notably, Ts-1, despite being a low-SD accession, consistently adopted the division strategy characteristic of high-SD accession Col-0.

### Accessions vary in their degree of coordination between sister division behaviors

The observation that accessions differ in their relative deployment of amplifying versus spacing divisions raised a fundamental question about coordination: do meristemoids and their sister SLGCs make division decisions independently, or are their behaviors coupled? Spacing divisions ensure stomatal separation by at least one intervening cell, a process requiring coordination among neighboring cells. Previous work has shown that such coordination regulates proliferative capacity between sister cells^32^ and among neighbors ^33^, generally by restricting stomatal production. We returned to our time course images to track the timing of SLGC spacing divisions relative to meristemoid fate decisions (division vs. differentiation). We referred to SLGCs that divided only after their sister meristemoid differentiated as “Type-I” divisions and SLGCs that divided while sister meristemoids were still undergoing amplifying division as “Type-II” divisions (Fig. 3A). Here, coordination refers to fate-coupling between sister cells rather than temporal synchrony. Low-SD accessions (Can-0, Kz-9, Ll-0) exhibited predominantly Type-I divisions, where SLGCs did not divide until their sister meristemoid differentiated, indicating stronger fate-based coordination between sister cells. In contrast, high-SD accessions (Col-0, Bur-0) showed weaker coordination, with only ~70% Type-I divisions (Fig. 3B). The prevalence of Type-II divisions in high-SD accessions is consistent with their higher stomatal production, as simultaneous proliferation of both sister cells generates more total meristemoids and SLGCs. Notably, Ts-1, despite being a low-SD accession, grouped with high-SD accessions in having reduced sister cell coordination. This indicates that Ts-1 achieves low stomatal production through a distinct developmental strategy from other low-SD accessions, revealing multiple developmental routes to similar phenotypic endpoints.

**Figure 3.**
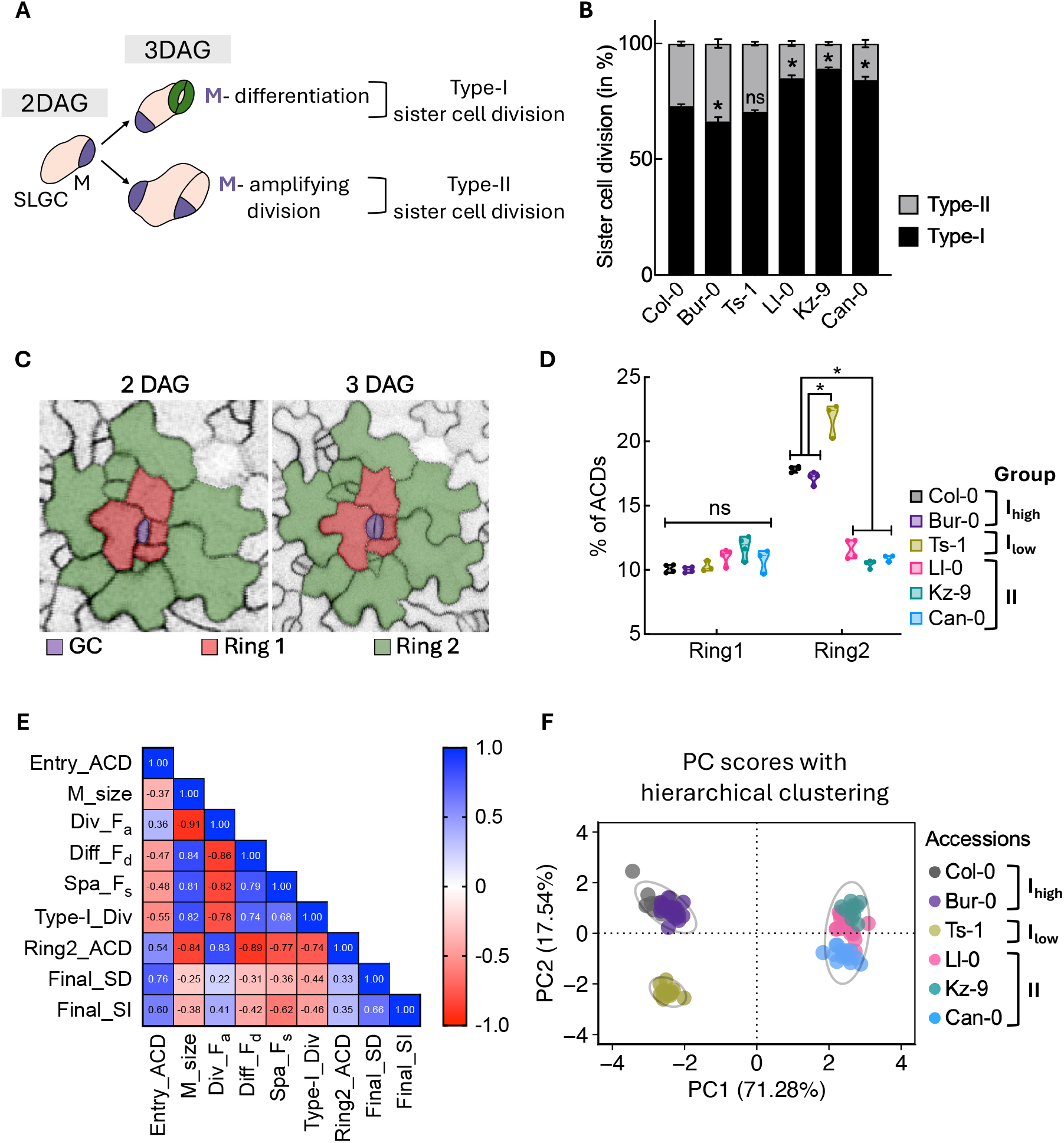
Natural variation in the impact of lineage relationships on stomatal division frequency. A. Schematic illustrating potential division behaviors among sister cells derived from an ACD; in Type I, the SLGC divides after the meristemoid differentiates, in Type II, both meristemoid and SLGC divide **B**. Ratio of division types in cells tracked in cotyledons from 2-3 DAG across accessions. * indicates difference from Col-0 ratio **C**. Micrographs of examples for tracing spatial organization of ACDs; For each differentiating GC pair (purple), immediate neighbors (red, Ring1) share a wall with the GCs. Ring 2 cells (green) contact Ring 1 cells, but not the purple GC. **D**. Number of cells undergoing ACDs in Ring1 and Ring2 in 24 time-course from 2-3 DAG. **E**. Pairwise Spearman correlations among all measured stomata lineage behaviors/outputs using pooled data from all accessions. Blue and red indicate positive and negative correlations, respectively. All correlations shown are statistically significant (p < 0.05, FDR corrected). **F**. Principal component analysis (PCA) of six Arabidopsis accessions on 9 measured variables (shown in E), each point represents an individual plant. Samples were grouped using hierarchical clustering (Ward’s method) on the first two principal components. Ellipses represent 95% confidence regions assuming multivariate normal distribution. Percentages of variance indicated on each PCA axis. Details on sample sizes and statistical tests are provided in Table S2.

Cell-cell signaling among non-sister neighbors is important for stomatal pattern ^34,35^. We therefore tested whether division coordination occurs only between sister cells or extends to local neighbors by tracing the fate of cells surrounding a differentiating GC. We classified cells that share a wall with the differentiating GC as ring 1, while cells located outside of these and sharing a wall with the ring 1 cells are classified as ring 2 (Fig 3C). In all six accessions, ring 1 cells underwent few cell divisions (Fig. 3C-D) consistent with previous observations that these immediate neighbors receive inhibitory signaling from GCs ^21^. In ring 2, low-SD accessions (Kz-9, Ll-0, Can-0) showed substantially fewer asymmetric divisions compared to high-SD accessions (Fig. 3C-D), reflecting an overall reduction in stomatal lineage cell division potential that limits SLGC-derived stomatal production. High-SD accessions (Col-0, Bur-0) along with Ts-1 exhibited higher overall division rates, with Ts-1 notably compensating for its lower entry divisions through the highest ring 2 division frequency (Fig. 1E, 3D). These three accessions shared a developmental strategy of lower meristemoid differentiation thresholds, biasing them toward increased meristemoid amplification over SLGC spacing division. Since each amplifying division generates an additional SLGC, this strategy inherently produces abundant SLGCs. Limiting subsequent SLGC spacing divisions prevents excessive stomatal overproduction. Conversely, low-SD accessions with reduced division rates employed larger meristemoid thresholds, favoring less amplification and more SLGC spacing divisions.

### Two distinct developmental strategies emerge from coordinated variation in cellular parameters

Our lineage tracing revealed that accessions differ across multiple cellular parameters: entry ACD frequency (Entry_ACD), meristemoid size threshold (M_size), meristemoid division (Div_F_a_), differentiation (Diff_F_d_), spacing division frequencies (Spa_F_s_), sister cell coordination (Type_I_div), and neighborhood proliferation (Ring2_ACD). Each parameter showed some association with final stomatal outcomes (Final_SD and Final_SI), but their relative importance and potential interdependencies remained unclear. To understand how these diverse cellular behaviors integrate to determine final stomatal outcomes, we performed comprehensive multivariate analyses. To quantify the contribution of these measured cell behaviors to overall stomatal production, we first performed pairwise Spearman correlation analysis among all the variables using bootstrapped, replicate-balanced datasets for each accession (Fig. 3E). While Entry_ACDs showed moderate positive correlations with Final_SD and Final_SI (r ~ 0.6-0.8), several other parameters like Spa_F_s_, Diff_F_d_ and Type_I_div exhibited comparable or even stronger associations with final stomatal traits indicating that post-entry developmental processes contribute significantly to stomatal patterning (Fig. 3E).

To further visualize how accessions differ across the multidimensional landscape of developmental parameters, we performed principal component analysis (PCA) incorporating all measured variables. The PCA clearly separated the accessions into distinct groups along the first two principal components, which together captured 88.82% of the total variance (Fig. 3F, S3A). PC1 divides Ts-1, Col-0 and Bur-0 (hereafter called Group I) from Can-0, Kz-9, and Ll-0 (Group II), notably placing low-SD accession Ts-1 with the high-SD accessions rather than with other low-SD accessions. PC2, which captures mostly final stomatal numbers, further distinguishes Ts-1 (Group I_low_) from Col-0 and Bur-0 (Group I_high_) (Fig. S3A). Thus, distinct combinations of developmental traits underlie the observed variation among accessions. These distinct, group-specific developmental strategies were further confirmed by separate subgroup correlation matrices (Fig. S3B-C) and partial least squares (PLS) regression analyses (Fig. S3D-F), which revealed that parameter importance varies substantially between accession groups. These patterns were obscured in the pooled correlation analysis where the opposing effects of the two regimes masked regime-specific relationships. Group I relied more heavily on meristemoid size and its consequent effects on division frequency and differentiation processes, while Group II depended more strongly on spacing divisions, and the coordination of divisions within sister lineages and neighboring cells (Fig. S3). The emergence of these two distinct developmental regimes from unbiased multivariate analysis suggests that cellular parameters do not vary randomly or independently across accessions. Instead, specific combinations of parameters co-occur, defining alternative developmental strategies for achieving stomatal pattern. Group I employs a dynamic, proliferation-intensive strategy with delayed meristemoid differentiation (due to a smaller size threshold), heavy reliance on meristemoid amplification, weak coordination between sister cells, and active neighborhood divisions. Group II employs a constrained, coordination-intensive strategy with early meristemoid differentiation (due to high size threshold), balanced contributions from meristemoids and SLGCs, strong sister coordination, and suppressed neighborhood activity.

### Stomatal lineage strategies dictate accessional responses to environmental fluctuations

Flexibility in stomatal lineage divisions has been proposed to promote environmental responsiveness and further, that particular developmental pathways might be sculpted by or for local environments ^19^. Among our accessions we found two major developmental regimes (Group I and Group II); thus, we could test whether this developmental diversity promotes or constrains the phenotypic plasticity of accessions in response to environmental changes.

To investigate response to temperature change, we germinated the selected accessions on agar plates under our standard light and temperature conditions before shifting them to high or low-temperature environments for 20 days (Material and Methods; see scheme in Fig S4A). Each accession displayed expected physiological responses to change in temperature, such as elongated hypocotyls and petioles in warmer conditions and compact rosette morphology in cooler conditions, suggesting that they had the appropriate machinery to sense and respond to temperature cues ^36^ (Fig. S4B-D). While all accessions decreased SD with increasing temperature, response magnitudes differed by regime. Group I_high_ showed the strongest responses, Group II the weakest, and Group I_low_ intermediate responses, indicating that developmental regime and environmental responsiveness are correlated but can be partially decoupled. Kz-9 was remarkably unresponsive, displaying stomatal density that remained nearly constant between 16°C and 28°C, with only the extreme 30°C condition eliciting a reduction (Fig 4A).

**Figure 4.**
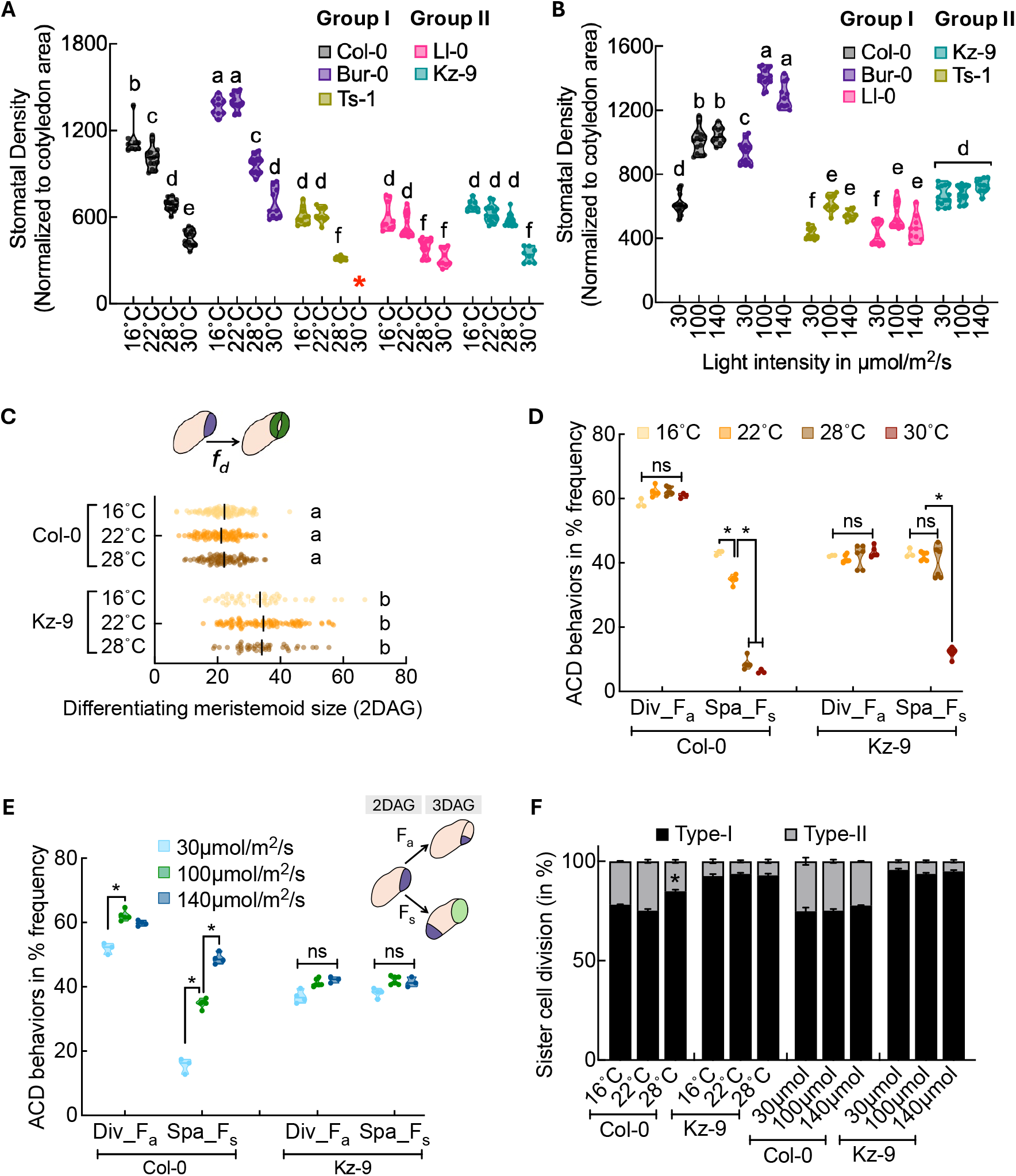
The capacity for developmental response to changes in temperature and light is conditioned by intra-lineage division coordination. **A-B** Violin plots showing stomatal density (20 DAG) in accessions grown at indicated temperatures **(A)** and light conditions **(B)**. Red asterisk in **(A)** represents the missing value of Ts-1 because seedlings could not survive at 30°C. Shape of violin indicates the data distribution density and different letters indicate significant difference (p < 0.01) **C**. Size-dependent differentiation of meristemoids (2DAG) is genotype, but not temperature specific as shown by the distribution of cell area at which meristemoids differentiate. Black vertical line indicates mean areal size of differentiating Ms, different letters indicate significant differences (p < 0.01). **D-E** Frequency of indicated division types (scheme in E) in accessions grown in indicated temperature (**D**) or light (**E**) conditions **F**. Ratio of division types in Col-0 and Kz-9 accessions under different temperature and light conditions from 2-3 DAG. Details on sample sizes and statistical tests are provided in Table S2.

In a similar experiment testing response to altered light intensity (Fig S4A-B, E) we observed the same trends in direction and magnitude of change, with Group I being more responsive than Group II and Kz-9 again being insensitive (Fig 4B). Correlating environmental response with developmental features, we noticed that Group I_high_ accessions in which meristemoids differentiate at a smaller size and exhibit less coordinated sister cell divisions show a stronger response to changes in temperature and light. In contrast, Group II accessions display a much weaker or completely dampened response to these environmental fluctuations. Thus, accessions with less coordinated divisions (Group I_high_: Col-0, Bur-0) exhibit higher plasticity, while more coordinated accessions (Group II: Ll-0, Kz-9) show developmental stability and reduced flexibility.

### Response to environmental change is primarily mediated by changes to spacing divisions

We next investigated precisely which parts of the stomatal lineage mediated environmental responses. In particular, because we showed several places where the lineage *could* be flexible, we were curious which ones would be used, and whether the same ones were used by different accessions and in response to different environmental inputs. For this analysis we focused on two accessions: Col-0, representing highly responsive Group I_high_ and Kz-9, the least responsive accession in Group II. We assayed features of the stomatal development pathway that defined the drivers of response to light and temperature change (Fig S4A).

We found that in both Col-0 and Kz-9 accessions, the meristemoid differentiates at a consistent size regardless of temperature conditions; a size that is closely linked to the meristemoid behavior, suggesting that the size threshold is genetically encoded and robust to external influences (Fig. 4C). Likewise, amplifying division frequency (Div_F_a_), which correlates with meristemoid size, did not change in either accession as we altered temperature (Fig. 4D). We did, however, observe a significant and directional change in spacing division frequency (Spa_F_s_) in the Col-0 accession under both altered temperature (Fig. 4D) and light conditions (Fig. 4E), with increased spacing divisions associated with higher SD and reduced spacing divisions correlating with lower SD. Spacing division frequency appears to be a major driver of stomatal density changes under these environmental cues, and it is abolished in Kz-9 except in the extreme case of 30°C (Fig. 4D-E). We also noted that the stringency of ACD sister-cell division correlation was sensitive to changing temperature in Col-0 but not in Kz-9 (Fig. 4F). Varying light conditions did not affect the sister cell division coordination in either Col-0 or Kz-9, however; thus different environmental cues can impact distinct aspects of the stomatal development pathway (Fig. 4F).

Taken together, these findings suggest that a decrease in stomatal density can arise through distinct developmental paths. Across the environmental conditions tested, changes in stomatal density were mainly by adjustments in spacing division frequency, as observed in the high stomatal density accession Col-0. In contrast, low stomatal density accession Kz-9 appeared to rely on a more fixed developmental strategy characterized by fewer entry ACDs, size-threshold– dependent meristemoid differentiation, and more coordinated spacing divisions. To integrate these parameter-specific responses into a comprehensive view of environmental plasticity, we performed PCA incorporating all developmental parameters measured across all temperature conditions tested on Col-0 and Kz-9 (Fig. 5A-B). Beyond accession-specific clustering along PC1 reflecting baseline developmental regimes, Col-0 and Kz-9 also separated along an environmental responsiveness axis (along PC2): Col-0 showed substantial displacement across environmental conditions, whereas Kz-9 remained nearly invariant, consistent with a constitutive developmental program exhibiting limited plasticity (Fig. 5A). The corresponding biplot indicates that this environmental axis is largely driven by final stomatal density and spacing division frequencies (Spa_F_s_), reaffirming that environmental responses target specific cellular parameters rather than globally altering the developmental program (Fig. 5B).

**Figure 5.**
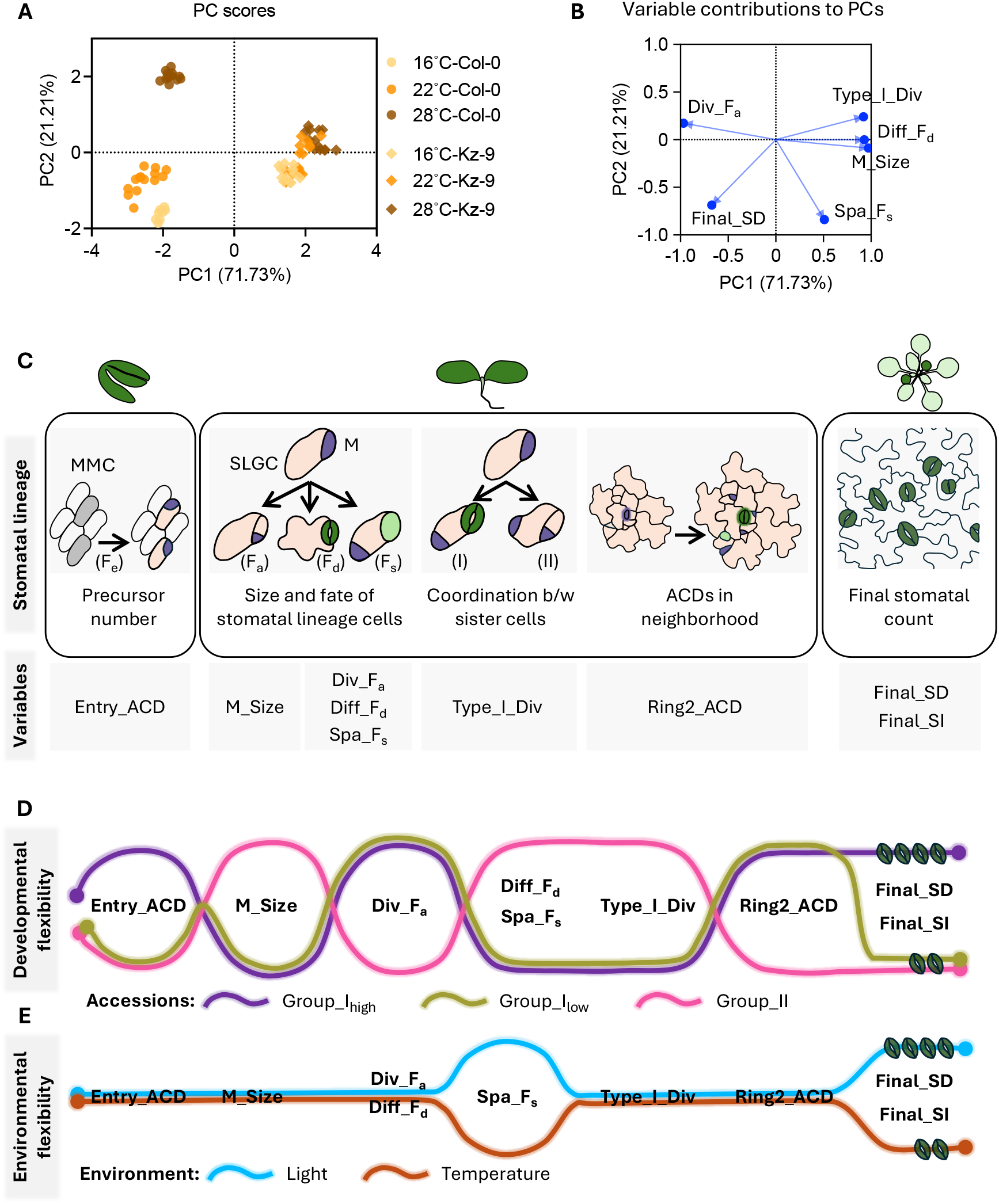
Model for flexibility modes in stomatal lineage. A. PCA plot of stomatal developmental parameters for Col-0 and Kz-9 grown at three temperatures (16°C, 22°C, 28°C), showing greater temperature-dependent dispersion in Col-0 along PC2. Each point represents a biological replicate under the indicated accession and temperature condition. **B**. Loading plot corresponding to panel **A** showing the contribution of individual developmental variables to the first two principal components (PC1 and PC2), **C**. Delineation of stomatal features from initiation through the stomatal lineage to final stomatal numbers with corresponding types of variation possible and observed at each stage noted below (variables). **D**. Graphical representation of two major regimes of stomatal lineage behaviors identified among accessions Group I_high_ is Col-0 and Bur-0; Group I_low_ is Ts-1; Group II is Ll-0, Kz-9 and Can-0. **E**. Graphical representation of lineage behavior regimes modulated to generate SI and SD changes in response to changing light and temperature. Details on sample sizes and statistical tests are provided in Table S2.

## DISCUSSION

Plasticity is a fundamental biological phenomenon that is increasingly important to understand as plants and animals encounter variable environmental conditions. The wide-spread existence of phenotypic plasticity is easy to document, but the genetic, cellular and developmental origins of such plasticity have been much harder to discover. Here we took a cellular and developmental view of plasticity in the generation of an adaptive trait: the density of stomata in plant leaves. By studying accessions with varying stomatal numbers, we were able to demonstrate their variability in cellular behavior and the coordination of division between sister cells and neighboring cells. Overall, the data support a model in which Arabidopsis accessions deploy divergent combinations of division strategies: size thresholds, division frequencies, and both sister cell and neighbor cell coordination to achieve their observed stomatal numbers (Fig. 5C-E). Within the diversity, two general regimes emerged from analysis of the six chosen accessions; group I accessions (Col-0, Bur-0, Ts-1) display a more cell-autonomous behavior where meristemoid cell size-based differentiation was a key factor, whereas Group II accessions (Ll-0, Kz-9, Can-0) were characterized by coordination of divisions among sister and neighbor cells (Fig. 5D). These two regimes appear to be convergent as accessions comprising group I or group II do not form close genetic or place of origin associations (Fig. S1A, H-I). We thus suggest that first, continuous stomatal variation emerges from distinct developmental programs rather than simple quantitative tuning. Second, that strategy does not determine the output; both Can-0 and Ts-1 reach low stomatal density but use alternative mechanisms. Third, parameters co-vary in coordinated patterns rather than independently, suggesting non-random relationships that may reflect shared genetic or developmental regulation.

Correlating environmental responsiveness to developmental features indicated that accessions with less coordinated divisions and smaller meristemoid size thresholds (Group I_high_) are more plastic, while those with more coordination and larger thresholds (Group II) are less plastic. Spacing division frequency emerged as the principal cellular process mediating environmental response (Fig. 5E). In Group I accessions, changes in SD under varying conditions stem almost entirely from altered rates of spacing divisions, whereas meristemoid size thresholds and amplifying division frequencies remain stable. The lack of spacing division flexibility in Group II accessions, especially Kz-9, emphasizes a genetically canalized pathway resistant to change. Recent studies show that vegetative phase change causes age-dependent increases in plasticity that vary across accessions ^37,38^, raising the intriguing question of whether the developmental regimes we identified shift during the juvenile-to-adult transition, and whether developmental architecture interacts with phase-specific regulatory programs to jointly shape environmental responsiveness.

Our goal in undertaking detailed characterization of the stomatal lineage in Arabidopsis accessions was to probe diversity in developmental trajectories and discern whether trajectories are functionally robust, converging on similar outcomes through different routes, and to what degree they may be flexible. The number of accessions we characterized is too small for genome-wide association studies, yet it is natural to wonder about the underlying genetic basis of the trajectories or developmental regimes we found. Genetic analysis of Arabidopsis herbarium specimens, for example, uncovered potential adaptive genetic variation in some of the non-essential stomatal regulatory genes ^39^. We therefore asked whether genes previously implicated in the stomatal lineage display any polymorphisms that suggest they underlie observed natural diversity. Although we identified missense mutations or indels in some key factors involved in stomatal division and fate (Fig. S5A), cell-cell signaling (Fig. S5B) and environmental response (Fig. S5C), none were consistently correlated with the developmental regimes we described in this work.

There is considerable interest in engineering plants with improved water use efficiency (WUE), CO_2_ capture, and stress tolerance ^40^. As valves regulating gas exchange between plants and the environment, and as the endpoint of the transpiration stream, stomata are attractive targets for such engineering ^41–43^. Natural or edited alleles affecting stomatal function or stomatal production can alter photosynthetic capacity or WUE but typically shift behavior in only one direction; for example, reducing stomatal numbers increases WUE, but can compromise performance under heat stress ^44,45^. Our findings suggest that rather than simply increasing or decreasing stomatal numbers, modulating the cellular parameters that govern developmental responsiveness could enable context-dependent optimization. In particular, SLGC spacing divisions emerged from our studies as a flexible lever for environmental adjustment (Fig 5E). No known stomatal regulator affects *only* this single parameter, but manipulations of the transcription factor SPEECHLESS or signaling peptides of the EPIDERMAL PATTERNING FACTOR family can affect spacing divisions ^18,46^. Applying precise engineering techniques to create novel synthetic expression patterns or tunable activity kinetics to target this specific part of the stomatal lineage may hold promise for generating plants that capitalize on developmental flexibility to become more climate resilient.

## ACKNOWLEDGEMENTS

We thank current and former Bergmann lab members for critical reading of manuscript. M.M. acknowledges financial support from the National Institute for Mathematics and Theory in Biology (Simons Foundation award MP-TMPS-00005320 and National Science Foundation award DMS-2235451). DCB is an investigator of the Howard Hughes Medical Institute. This research was supported in part by Grants No. NSF PHY-1748958 and No. NSF PHY-2309135 and the Gordon and Betty Moore Foundation Grant No. 2919.02 to the Kavli Institute for Theoretical Physics.

## AUTHOR CONTRIBUTIONS

MR, conceptualization, data generation, data analysis, quantification, statistical analysis, data visualization, writing; MM, statistical analysis, data visualization; NS, data generation; DCB, conceptualization, supervision, funding, writing and editing

## DATA AVAILABILITY

All data needed to evaluate the conclusions in the paper are present in the paper, and raw data are available at doi:10.6084/m9.figshare.32011776. This study did not generate unique code.

## COMPETING INTERESTS

The authors declare no competing interests.

## SUPPLEMENTAL INFORMATION

**STAR Methods**

**Key resources table**

**Figures S1-S5** and supplemental references

**Table S1:** Origin of accessions and associated climate data,

**Table S2:** Statistical information about all data presented in this paper

## Notes

https://doi.org/10.6084/m9.figshare.32011776

### Competing Interest Statement

The authors have declared no competing interest.

### Summary of Updates

We have clarified statistical approaches, strengthened our discussion of developmental diversity across dicots, toned down speculative evolutionary interpretations, improved figure clarity and consistency, and enhanced the detail and transparency of the methods section.

