## Supplementary material for "Stomatal setpoints and environmental responsiveness are sculpted by developmental trajectories": Methods; Supplemental Figures 1-5; Tables 1-2

**This file contains**
Star Methods and Key resources Table
Supplemental Figures S1-S5
Supplemental Tables S1-S2
Supplemental References

**Raw data** are available at doi:10.6084/m9.figshare.32011776

### STAR METHODS

#### Experimental model and subject details

Seeds of *Arabidopsis* accessions (listed in Key Resources Table and Supplementary Table S1) were surface-sterilized using either ethanol or chlorine gas, then stratified for three days in dark at 4˚C. Following stratification, seeds were grown in a Percival growth chamber (CU36L5) on half-strength Murashige and Skoog (MS) medium solidified with 0.8% agar, under long-day conditions (16 h light / 8 h dark at 22 °C) with moderate-intensity, full-spectrum light (110 μE) as "standard conditions". The age of the seedlings is indicated as DAG (days after germination), which was standardized and synchronized across accessions based on radicle protrusion. All accessions germinated synchronously except Ll-0, which lagged by ~24 h.

#### Method details

*Arabidopsis transformations*

To generate transgenic marker lines in Arabidopsis accessions Col-0, Bur-0, Kz-9, Ll-0, Ts-1, and Can-0, the *pATML1::mCherry-RCI2A* construct (Key resources table) was introduced into *Agrobacterium tumefaciens* and transformed into plants via the floral dip method [47]. Transgenic seedlings were selected on ½ MS medium containing 50 μg/mL kanamycin. All experiments were performed using homozygous T3 lines.

*Environmental conditions*

For light and temperature experiments, all accessions were initially grown under standard conditions (110 μE light, 22 °C) for 24 hours to induce germination. They were then shifted to either high light (140 μE) or low light (30 μE), and high temperature (28 °C and 30˚C) or low temperature (16 °C) for 3 days (for the 24-hour time-course analysis) or 22 days (for final guard cell counts). This experimental design ensured that the initial entry ACD–derived precursor pool, established within the first 24-hour, was standardized across environmental treatments.

*Phenotypic characterization*

Images of hypocotyls (vertically grown), cotyledons, and full rosettes were taken at defined time points after germination on ½ MS medium under specified growth conditions, as indicated in the figure legends. For quantification of mature stomatal phenotypes, seedlings of the specified ages were cleared overnight in a 7:1 ethanol:acetic acid solution, rinsed in 1 M KOH for 1 h, and mounted in Hoyer's medium (7.5 g gum arabic, 5 mL glycerol, 100 g chloral hydrate, 30 mL $H_2O$). Differential interference contrast (DIC) images were acquired using a Leica DMi7 microscope from two regions on either side of the midvein of the 21 DAG abaxial cotyledon epidermis, unless stated otherwise. All stomata within the imaged sectors were manually counted, and the cotyledon area was measured using the CellCounter plugin in Fiji (ImageJ)[48]. Stomatal density (SD) was calculated as the total number of stomata per unit area, stomatal index (SI) as the number of stomata relative to the total number of epidermal cells, and the stomatal number/density per cotyledon by normalizing SD to the total cotyledon area in 20 DAG cotyledons.

*Microscopy, image acquisition and image analysis*

All fluorescence imaging and time-course (lineage tracing) experiments were performed on a Leica Stellaris 8 confocal microscope with HyD detectors using 25x or 40x NA1.1 water objective with image size 1024 X 1024 pixels and a digital zoom from 1x to 2x. Raw Z-stacks were projected with Sum Slices in Fiji[48]. Entry ACDs were quantified at 1 DAG, when cotyledons are just opening, by counting physically asymmetric sister-cell pairs.

To track cell fates, 2 DAG seedlings expressing the *pATML1::mCherry-RCI2A* plasma membrane marker were mounted in water on slides sealed with vacuum grease with sufficient space to avoid compressing tissues, imaged, and returned to ½ MS plates under specified light and temperature conditions. The same seedlings were re-imaged after 24 and 48 hours to capture the developmental outcomes of stomatal lineage cells in the epidermis. Meristemoid size measurements were taken from the abaxial cotyledons at 2 DAG. Differentiating and dividing meristemoids were categorized based on their behavior recorded after 24 hours (division/ACD vs. differentiation). The protocol does not allow us to record the exact meristemoid size at birth, however, using this time course method, the size threshold for meristemoids in Col-0, the reference accession, was comparable to previously reported values[17].

The frequencies of meristemoid division (amplification division), SLGC division (spacing division), and meristemoid differentiation were calculated from the 24-hour lineage-tracing time-course images, and the number of amplification divisions undergone by the meristemoid before differentiation (e.g. in Fig. 2B) was determined using the additional 48-hour time point.

To assess sister cell division correlations across accessions, asymmetrically divided sister cell pairs were tracked for 24 hours (2–3 DAG). The frequency at which both cells divided together (type II division) or where the larger SLGC divided only after the meristemoid committed to differentiate via symmetric division (type I division) was calculated. Neighbor coordination was measured from the same time-course images by tracking asymmetric divisions (ACDs) surrounding differentiating guard cells (GCs) during 2–3 DAG. Ring 1 cells were defined as immediate neighbors sharing a border with the GC, and ring 2 cells as the next layer of neighbors bordering ring 1. The frequency of ACDs occurring within ring 1 and ring 2 cells was then plotted.

*Hoechst staining and nuclear area measurement*
For nuclear area measurements, Hoechst-stained nuclei of 3 DAG cotyledons[49] were imaged by confocal microscopy, and nuclei size of the meristemoid (based on cell morphology and size using plasma membrane marker lines) were manually measured using ImageJ/Fiji.

*SNPs analysis and mapping*
Single nucleotide polymorphism (SNP) variants in selected genes were obtained from the POLYMORPH 1001 tool of the 1001 Genomes Project[50] for all chosen accessions. Corresponding in-frame insertions/deletions and missense variants causing amino acid changes were mapped onto the Col-0 reference protein sequence using Geneious Prime software. The analysis specifically focused on SNPs located within the coding regions of the selected genes.

*Accession numbers*
TAIR accession number of the genes used in Fig. S5 are: SPCH (AT5G53210), MUTE (AT3G06120), FAMA (AT3G24140), SCRM (AT3G26744), SCRM2 (AT1G12860), HDG2 (AT1G05230), EPFL9 (AT4G12970), EPF1 (AT2G20875), EPF2 (AT1G34245), SDD1 (AT1G04110), TMM (AT1G80080), ERECTA (AT2G26330), ERL1 (AT5G62230), PIF4 (AT2G43010), PIF5 (AT3G59060), PHYB (AT2G18790)

**Climate data**
Climatic information for the Arabidopsis accessions was obtained from AraCLIM V2.0[51,52] (https://gramene.org/CLIMtools/arabidopsis_v2.0/AraCLIM-V2/)**,** an online resource that integrates high-resolution environmental data for natural *Arabidopsis thaliana* accessions from the 1001 Genomes Project[50]. For consistency, only variables derived from **CHELSA** (*Climatologies at High Resolution for the Earth's Land Surface Areas*) were used.

**Quantification and statistical analysis**
All statistical analysis in this study were performed using GraphPad Prism (v. 10.5.0) and RStudio (v. 2025.05.1+513 "Mariposa Orchid"). Pairwise comparisons between accessions or conditions were performed using either an unpaired Student's *t*-test (for sample sizes >10, assuming normal distribution) or a Mann–Whitney U test (for sample sizes ≤10 or non-normal distributions). For datasets involving multiple groups, one-way ANOVA was performed followed by Tukey's Honest Significant Difference (HSD) post-hoc test for pairwise comparisons. $p < 0.01$ was used as the significance cut-off. For all graphs, ns- not significant, * - $p < 0.01$ and different individual letter indicate group of samples with statistically significant difference between their means (BKY corrected $p$ value < 0.01). Details of statistical tests, including methods used, number of replicates, definitions of mean and error bars, and significance values for individual graphs, are provided in Table "Statistical Information, Rath et al" (Table S2).
**Simple linear regression (**Ordinary least square) was performed to examine relationships between entry events and stomatal traits (stomatal density and stomatal index) across *Arabidopsis* accessions using the lm() function in R. Models were fitted either with all accessions combined or separately for each accession. Replicate numbers within each accession were balanced using a stratified nonparametric bootstrap, resampling trait values with replacement via the sample() function in base R within a tidyverse (*dplyr*, *tidyr*) framework. The resulting balanced datasets were used to assess and visualize rank-based associations across accessions. The coefficient of determination (**$R^2$**) was extracted to estimate the variance explained by entry events, and residuals were calculated and plotted against entry events to evaluate model fit. Accession-level Best Linear Unbiased Estimators (BLUEs) were estimated using mixed models fitted with the lmer() function from lme4 package in R, with replicates treated as random effects, and adjusted means extracted using the emmeans() package. This approach generates a single adjusted mean per accession that accounts for environmental variation and measurement error across replicates. All regression analyses and visualizations were performed in RStudio using base R and ggplot2.

**Pairwise non-parametric Spearman rank correlation** coefficients were computed among all the developmental parameters (variables) using the bootstrapped resampled data from all accessions combined, group I accessions (Col-0, Bur-0 and Ts-1) and group II accessions (Ll-0, Kz-9 and Can-0) separately. Correlation coefficients (ρ) were visualized as color-coded heatmaps using GraphPad Prism.
**Partial least square (PLS) regression** model was constructed using the plsr() function from pls package in R, with the final stomatal index as the response variable and all developmental variables as predictors. Variable Importance in Projection (VIP) scores were calculated to quantify the contribution of each predictor variable to the PLS model and variables with VIP > 1 were considered highly influential.
**Principal Component Analyses (PCA)** were conducted using prcomp() on bootstrapped and resampled data to visualize multivariate structure: (1) nine developmental parameters across six accessions, (2) six developmental plus environmental parameters (temperature) for Col-0 and Kz-9 under 16°C, 22°C, and 28°C, (3) climatic variables (latitude, longitude, mean temperature, precipitation, and seasonality) from the **AraCLIM V2.0 online tool** for nine accessions, and (4) pairwise genetic distances among Arabidopsis accessions as mentioned in Xu et al.,2025[53]. To determine the genetic distance, we used the genetic variation from the 1001 Genomes Project (1001genomes_snp-short-indel_only_ACGTN.vcf.gz from the https://1001genomes.org/data/GMI-MPI/releases/v3.1/ website [accessed 9/10/2025]) and computed the p-distance matrix with VCF2Dis v1.55 [50,53]. PC scores were extracted and visualized in a biplot showing the first two principal components (PC1 and PC2), with data points colored by accession. Hierarchical clustering of PCA scores was performed using hclust(method = "ward.D2") on Euclidean distances computed from PC1 and PC2; clusters were defined using cutree(k = 3) and visualized as 95% confidence ellipses in ggplot2. All visualizations were generated using ggplot2 and assembled in Microsoft PowerPoint.

### KEY RESOURCES TABLE

| REAGENT or RESOURCE | SOURCE | IDENTIFIER |
|---|---|---|
| **Bacterial and virus strains** | | |
| *Escherichia coli:* strain TOP10 | N/A | N/A |
| *Agrobacterium tumefaciens* GV3101 RifR, GentR with pSOUP plasmid (TetR) | N/A | N/A |
| **Chemicals, peptides, and recombinant proteins** | | |
| Hoechst 33342, trihydrochloride trihydrate | Life Technologies | H3570 |
| FM 4-64 Dye | Thermo Fisher Scientific | T3166 |
| MS basal salt | Caisson Laboratories (now Plant Cell Technology, PCT) | MSP01-50LT |
| MES buffer | Caisson Laboratories (now Plant Cell Technology, PCT) | M009-500GM |
| Micropropagation Agar-Type 1 | Caisson Laboratories (now Plant Cell Technology, PCT) | A038 |
| **Experimental models: Organisms/strains** | | |
| *Arabidopsis thaliana* accession Can-0 | ABRC (https://abrc.osu.edu/) | CS6660 |
| *Arabidopsis thaliana* accession Ts-1 | ABRC (https://abrc.osu.edu/) | CS22647 |
| *Arabidopsis thaliana* accession Sp-0 | ABRC (https://abrc.osu.edu/) | CS76603 |
| *Arabidopsis thaliana* accession C-24 | ABRC (https://abrc.osu.edu/) | CS22680 |
| *Arabidopsis thaliana* accession Jm-0 | ABRC (https://abrc.osu.edu/) | CS76520 |
| *Arabidopsis thaliana* accession Shahdara | ABRC (https://abrc.osu.edu/) | CS22652 |
| *Arabidopsis thaliana* accession Ll-0 | ABRC (https://abrc.osu.edu/) | CS77047 |
| *Arabidopsis thaliana* accession Kz-9 | ABRC (https://abrc.osu.edu/) | CS76537 |
| *Arabidopsis thaliana* accession Col-0 | ABRC (https://abrc.osu.edu/) | CS76778 |
| *Arabidopsis thaliana* accession Bur-0 | ABRC (https://abrc.osu.edu/) | CS76734 |
| *Arabidopsis thaliana* accession Van-0 (harbors a natural loss of function allele in *ERECTA*, a known stomatal pattern regulator) | ABRC (https://abrc.osu.edu/) | CS22694 |
| *Arabidopsis* Col-0: WT pAtML1::mCherry-RCI2A | Davies and Bergmann 2014[47] | N/A |
| *Arabidopsis* Bur-0: WT pAtML1::mCherry-RCI2A | This paper | N/A |
| *Arabidopsis* Ll-0: WT pAtML1::mCherry-RCI2A | This paper | N/A |
| *Arabidopsis* Kz-9: WT pAtML1::mCherry-RCI2A | This paper | N/A |
| *Arabidopsis* Ts-1: WT pAtML1::mCherry-RCI2A | This paper | N/A |
| *Arabidopsis* Can-0: WT pAtML1::mCherry-RCI2A | This paper | N/A |
| **Recombinant DNA** | | |
| R4pGWB401 pAtML1::mCherry-RCI2A | Davies and Bergmann 2014[47] | |
| **Software and algorithms** | | |
| ImageJ/FIJI version 2.14.0/1.54f | Schindelin et al.,2012[49] | https://imagej.net/software/fiji/ |
| GraphPad Prism 10.5.0 | Dotmatics, Boston, MA | https://www.graphpad.com/features |
| Geneious Prime 2020.0.5 | Dotmatics, Boston, MA | https://www.geneious.com/ |
| Microsoft Office 365 version 16.97.2 (25052611) | Microsoft, Redmond, USA | N/A |

| | | |
|---|---|---|
| Leica Application Suite X | Leica Microsystems; Mannheim, Germany | N/A |
| RStudio | Posit Software, PBC | https://posit.co/download/rstudio-desktop/ |
| R (v. 2025.05.1+513 “Mariposa Orchid”) | R Core Team | https://www.r-project.org/ |
| **Other** | | |
| Confocal microscope Stellaris | Leica Microsystems; Mannheim, Germany | |
| Arabidopsis Information Resource (TAIR) | N/A | https://www.arabidopsis.org/ |
| AraCLIM V2.0 | Ferrero-Serrano et al., 2022[52] Ferrero-Serrano A and Assmann SM, 2019[53] | https://gramene.org/CLIMtools/arabidopsis_v2.0/AraCLIM-V2/ |

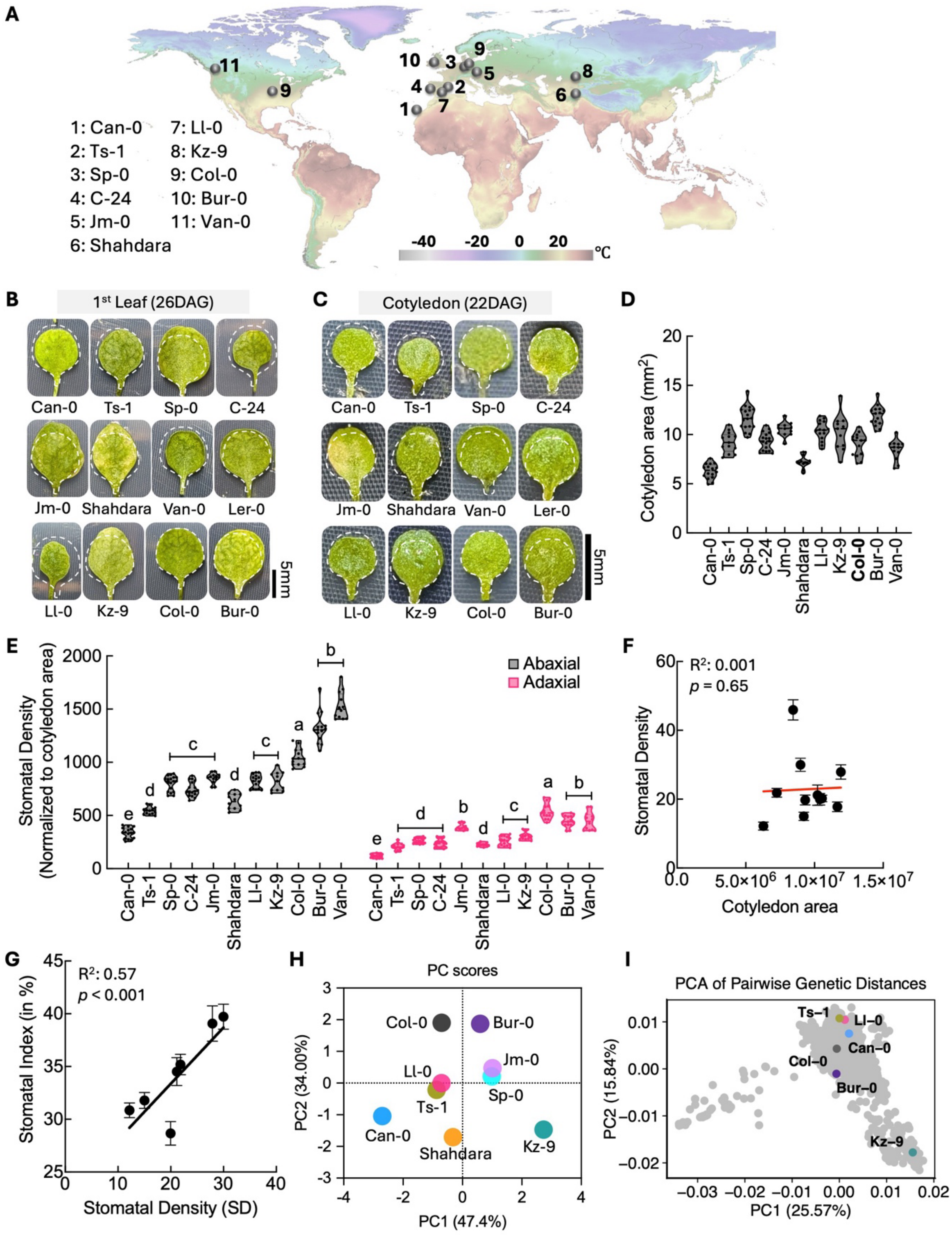

**Figure S1 Additional characterization of accessions used in this study A.** Place of origin of accessions, with average yearly temperature indicated by color coding, map adapted from www.worldclim.org **B-C.** Representative images of leaves (**B**) and cotyledons (**C**) from the indicated Arabidopsis accessions grown under standard long day conditions (16 h light / 8 h dark at 22 °C with full-spectrum light, 110 μE). White dotted outline is taken from Col-0 and overlaid on accessions to illustrate relative differences in overall size and shape. **D.** Cotyledon area measured at 22 DAG **E.** Stomatal density (SD) calculated per cotyledon area on abaxial (black) and adaxial (pink) sides of cotyledons at 20 days after germination (DAG) **F**. Linear regression of cotyledon area and SD showing no correlation between these parameters across 11 measured accessions. **G**. Linear regression of SD and Stomatal Index (SI) showing a positive correlation. **H.** Principal component analysis (PCA) of nine Arabidopsis accessions based on five environmental variables (latitude, longitude, annual mean temperature, annual mean precipitation, and temperature seasonality) from their native collection sites. Environmental data obtained from AraCLIM V2.0. Accessions do not cluster based on their climatic regions of origin. PC1 and PC2 explain 47.4% and 34.00% of the variance, respectively. **I.** PCA of 1,135 accessions (grey) from the 1001 Genomes Project [1] based on pairwise genetic distances [2], with the six accessions selected for this study highlighted in color. PC1 and PC2 explain 25.57% and 15.84% of the variance, respectively.

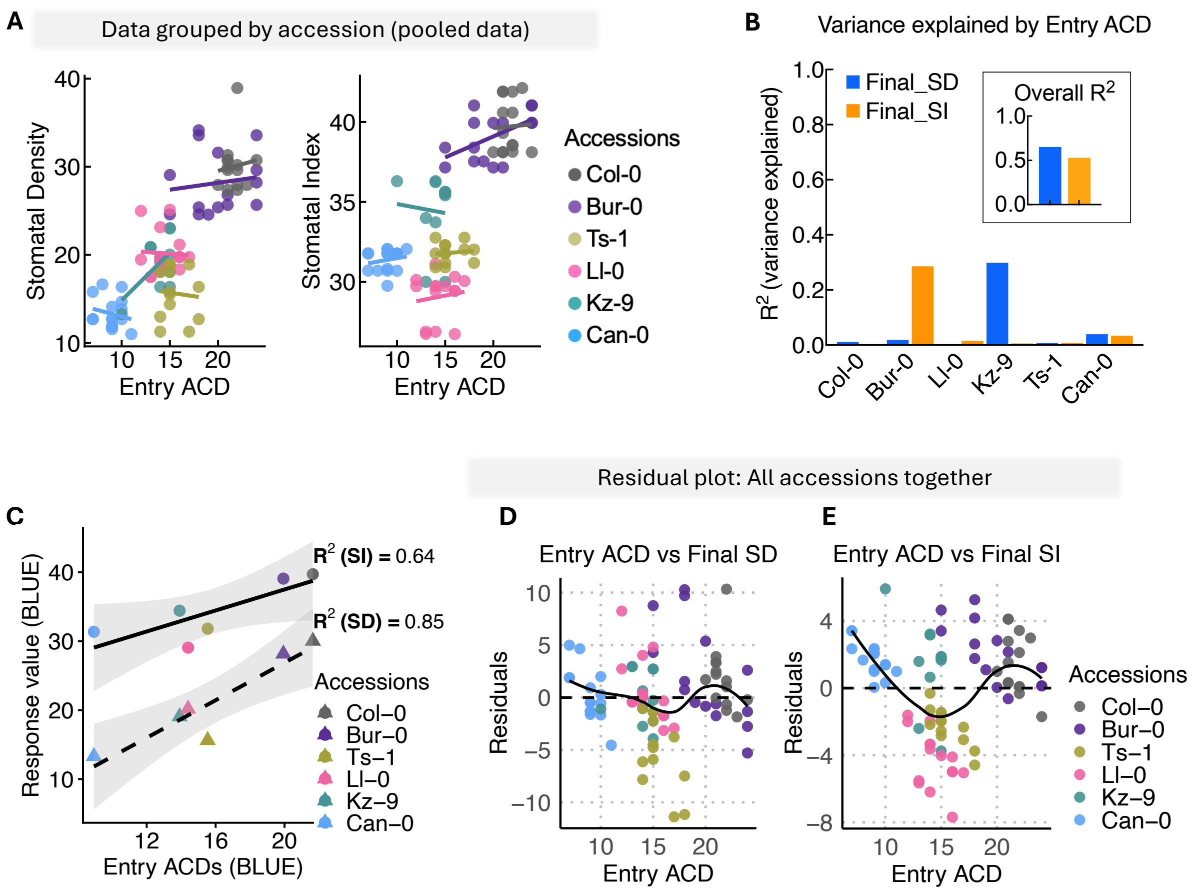

**Figure S2 Entry events show limited predictive power for final stomatal traits, suggesting additional developmental factors**

**A.** Simple linear regression with data grouped by accession, analyzing stomatal density (left) and stomatal index (right) separately as response variables. **B**. $R^2$ values from ordinary least squares (OLS) regression between entry events and stomatal density (SD) or stomatal index (SI), shown for all accessions combined (overall) and individually (from Fig. 1F and S2A). Blue and orange bars represent SD and SI models, respectively. **C.** Simple linear regression of genotype-adjusted means (BLUEs) from mixed models accounting for biological replicates. Dots are color coded by accession with each dot representing one accession's BLUE for entry ACDs versus final SD (triangles, dashed line) or SI (circles, solid line). $R^2$ values shown in the plot **D-E**. Residual plot for the pooled linear regression model in panel B, displaying residuals (observed minus predicted values) against entry events for final SD **(D)** and final SI **(E)**. The non-random distribution (U-shaped curve) and accession-specific clustering indicate that entry events alone are insufficient to fully explain variation in final stomatal index, suggesting contributions from additional factors or genotype-specific effects.

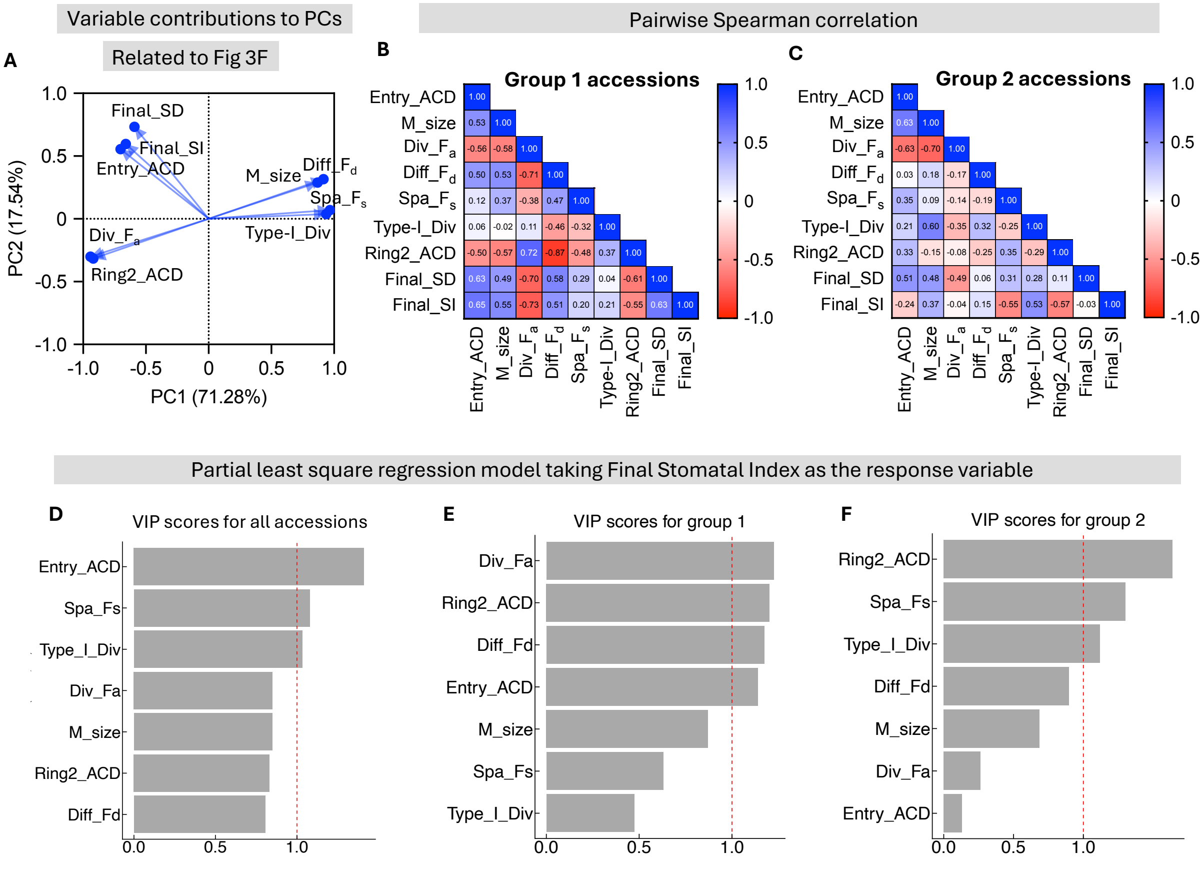

**Figure S3 Pairwise correlations and variable importance of stomatal developmental parameters across accessions.**

**A.** Loading plot corresponding to Fig. 3F showing the contribution of individual developmental variables to the first two principal components (PC1 and PC2). **B-C** Pairwise Spearman correlation matrices showing relationships among stomatal developmental parameters for Group 1 accessions (Col-0, Bur-0, and Ts-1) **(B)** and Group 2 accessions (Ll-0, Kz-9, and Can-0) **(C)**. Color intensity and numerical values indicate correlation strength, with blue representing positive correlations and red representing negative correlations. **D-F** Variable Importance in Projection (VIP) scores from partial least square regression models with final stomatal index as the response variable. VIP scores indicate the contribution of each developmental parameter to predicting the final stomatal index for all accessions combined **(D)**, Group 1 accessions only **(E)**, and Group 2 accessions only **(F)**. The red dashed line indicates the threshold for significant importance (VIP > 1.0). Higher VIP scores indicate greater contribution to the model's predictive ability.

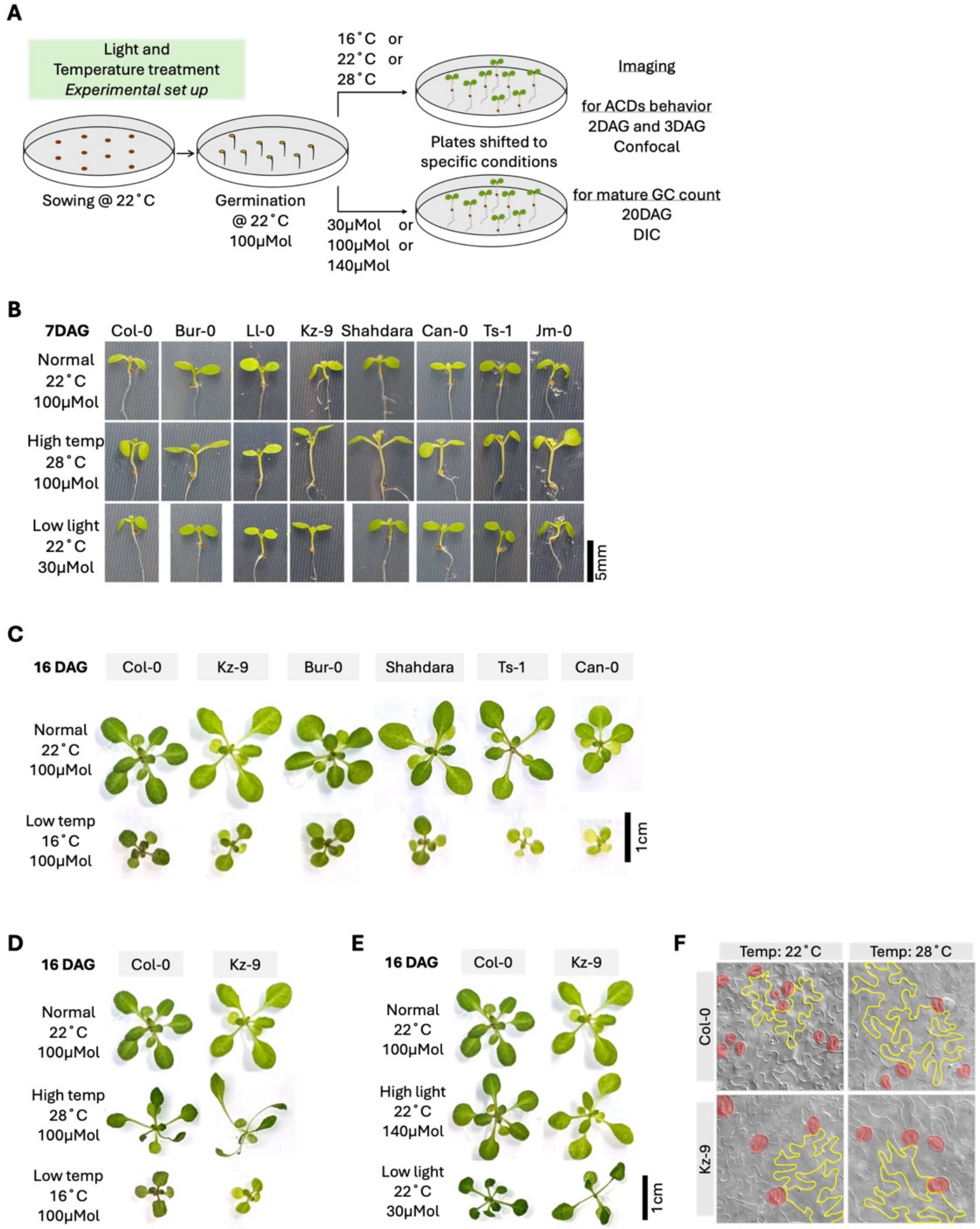

**Figure S4 Experimental setup for temperature and light perturbations and additional, non-stomatal phenotypes under these conditions (related to Figure 4)**

**A**. Schematic of environmental change experiments **B**. Behavior of seedlings exposed to altered environments. All accessions displayed the hypocotyl elongation and cotyledon expansion phenotypes expected from previous studies[3]. **C.** Comparison of whole plant responses at 16 DAG in other ecotypes. D-E**.** Comparison of whole plant responses to three different temperature (D) and light regimes (E) in Col-0 and Kz-9 at 16 DAG, indicate that Kz-9 is capable of sensing and responding in terms of petiole extension and rosette morphology, even if it does not exhibit stomatal lineage change. **F.** Micrographs of epidermis in cleared cotyledon tissue of Col-0 and Kz-9 showing that epidermal cell morphology (yellow highlight) of Kz-9 at 22˚C resembles Col-0 at 28˚C (Stomata are false colored red).

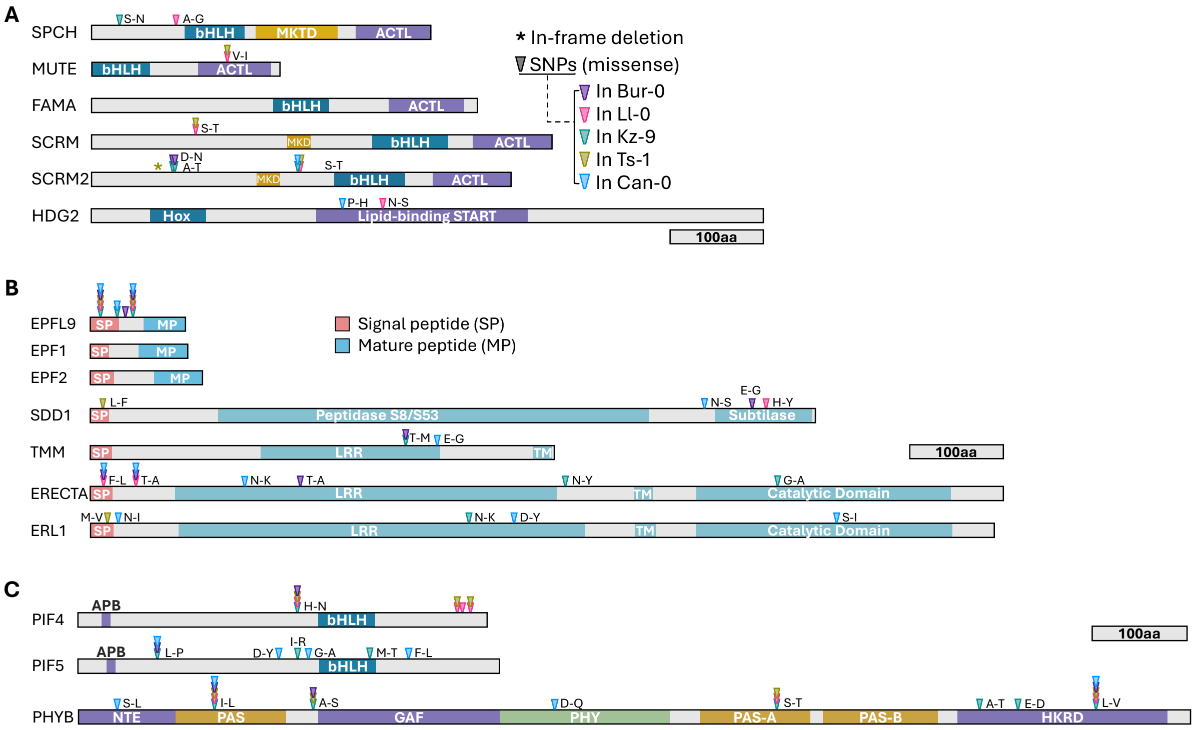

**Figure S5 Variation in known stomatal regulation genes does not correlate with differences in stomatal lineage behaviors among accessions A-C.** Protein structure diagrams showing single nucleotide polymorphisms (SNPs) in key stomatal developmental genes across selected accessions. **(A)** Transcription factor genes **(B)** Stomatal cell-cell communication genes **(C)** Light and temperature perception/response genes. SNP locations are marked with colored arrows indicating the specific accession. Although SNPs are present in some accessions, none correlate with the different division patterns, SI, SD or ability to respond to light or temperature seen across the measured accessions.

**Table S1: Geographic origin and environmental conditions of accessions analyzed in this work.**

| *Arabidopsis thaliana* Accession name | TAIR ID | Geographic coordinates Longitude | Latitude | Location of origin | Annual mean temperature |
|---|---|---|---|---|---|
| Can-0 | CS6660 | -13.4811 | 29.2144 | Las Palmas, Spain | 18.35974503 |
| Ts-1 | CS22647 | 2.93056 | 41.7194 | Tossa de Mar, Spain | 16.31507301 |
| Sp-0 | CS76603 | 13.181 | 52.5339 | Berlin, GERMANY | 9.557936668 |
| C-24 | CS22680 | -8.45 | 41.25 | Seroa, Portugal | 16.16753387 |
| Jm-0 | CS76520 | 15 | 49 | Czech Republic | 8.66870594 |
| Shahdara | CS22652 | 68.48 | 38.35 | Shakhrinaw, Tajikistan | 16.33777618 |
| Ll-0 | CS77047 | 2.49 | 41.59 | Spain | 15.36473656 |
| Kz-9 | CS76537 | 73.1 | 49.5 | Kazakhstan | 3.402762175 |
| Col-0 | CS76778 | -92.3 | 38.3 | Missouri, US | 13.6827879 |
| Bur-0 | CS76734 | -6.2 | 54.1 | Ireland | 10.11813259 |
| Van-0 | CS22694 | -123.206 | 49.2655 | Canada | 10.32006454 |

**Table S2: Statistical information for all data in Rath et al, 2026**

| Figure panel | Description of the data | Sample size | Error bars | Statistical test |
|---|---|---|---|---|
| 1B | Stomatal Density in 20 DAG abaxial cotyledon across accessions | 14-18 individual cotyledons per accession | Median | ANOVA with post-hoc Tukey's HSD test |
| 1C | Stomatal Index in 20 DAG abaxial cotyledon across accessions | 6-11 individual cotyledons per accession | Median | ANOVA with post-hoc Tukey's HSD test |
| 1E | Number of entry ACDs and fully formed GC in 1 DAG abaxial cotyledon across accessions | 08-12 individual cotyledons per accession | Median | ANOVA with post-hoc Tukey's HSD test |
| 1F | Simple linear regression of entry ACD vs final SD and final SI | bootstrapped, replicate-balanced datasets (n ≈ 14–15 per accession) across six accessions | Shaded region- 95% CI | Ordinary Least Square (OLS) regression analysis |
| 1G | Comparison of entry ACDs, final stomatal outcomes in percentage across accession | 08-11 individual cotyledons per accession | Standard deviation | Mann-Whitney test |
| 2A | Size of meristemoid that undergoes differentiation or ACD in 2 DAG abaxial cotyledon across accessions | Combined 30-40 cells per each of 3 different individuals | Shaded region- transition size | Logistic regression between ACD and Diff, ANOVA with post-hoc Tukey's HSD test (for the transition size) |
| 2B | Nuclear area of meristemoids in 3 DAG abaxial cotyledon across accessions | Combined 30-40 cells per each of 3 different individual cotyledons | Median | ANOVA with post-hoc Tukey's HSD test |
| 2C | Rounds of amplifying divisions undergone in 2-4 DAG abaxial cotyledon across accessions | 60 cells per each of 3 different individual cotyledons | 95% CI | Mann-Whitney tests |
| 2D | Meristemoid behavior in 2-3 DAG abaxial cotyledon across accessions | 60 cells per each of 4-7 different individual cotyledons | Median | Mann-Whitney tests |
| 2E | SLGC division frequency (%) in 2-3 DAG abaxial cotyledon across accessions | 60 cells per each of 4-7 different individual cotyledons | Median | Mann-Whitney tests |

| | | | | |
|---|---|---|---|---|
| 3B | Sister cell division (in %) in 2-3 DAG abaxial cotyledon across accessions | 25-30 cells per each of 3-6 different individual cotyledons | SEM | Mann-Whitney tests |
| 3D | Percentage of ACDs around a stoma in 2-3 DAG abaxial cotyledon across accessions | 60 stomata per each of 3-5 different individual cotyledons | Median | Unpaired t tests |
| 3E | Pairwise correlation among stomatal development variables | bootstrapped, replicate-balanced datasets (n ≈ 14–15 per accession) across six accessions | N/A | Spearman's rank correlation |
| 3F | PCA plot of 6 accessions on 9 measured variables | bootstrapped, replicate-balanced datasets (n ≈ 14–15 per accession) across six accessions | Grey ellipse-95% CI | Principal Component Analysis with Hierarchical clustering (Ward's method, Euclidean distance) |
| 4A | Stomatal Density in 20 DAG abaxial cotyledon across accessions under different temperature conditions | 14-18 individual cotyledons per accession | Median | ANOVA with post-hoc Tukey's HSD test |
| 4B | Stomatal Density in 20 DAG abaxial cotyledon across accessions under different light conditions | 14-18 individual cotyledons per accession | Median | ANOVA with post-hoc Tukey's HSD test |
| 4C | Meristemoid behavior in 2-3 DAG abaxial cotyledon across accessions under different temperature conditions | 60 cells per each of 3 different individual cotyledons per accession | Mean with 95% CI | ANOVA with post-hoc Tukey's HSD test |
| 4D | ACD behavior frequency (in %) in 2-3 DAG abaxial cotyledon across accessions under different temperature conditions | 60 cells per each of 4-6 different individual cotyledons | Median | Mann-Whitney test |
| 4E | ACD behavior frequency (in %) in 2-3 DAG abaxial cotyledon across accessions under different light conditions | 60 cells per each of 4-6 different individual cotyledons | Median | Mann-Whitney test |
| 4F | Sister cell division (in %) in 2-3 DAG abaxial cotyledon across accessions under different temperature and light conditions | ~30 cells per each of 3-5 different individual cotyledons | SEM | Mann-Whitney test |
| 5A | PCA plot of Col-0 and Kz-9 on 6 measured variables plus temperature as environmental variable | bootstrapped, replicate-balanced datasets (n ≈ 10-14 per accession) of Col-0 and Kz-9 under mentioned conditions | N/A | Principal Component Analysis |
| 5B | Loading plot of the PCA plot | bootstrapped, replicate-balanced datasets (n ≈ 10–14 per accession) of Col-0 and Kz-9 under mentioned conditions | N/A | N/A |
| S1D | Cotyledon area of the indicated accession at 22DAG | 14-18 individual cotyledons per accession | Median | N/A |
| S1E | Stomatal Density normalized to cotyledon area in 20 DAG abaxial and adaxial cotyledon across accessions | 14-18 individual cotyledons per accession | Median | ANOVA with post-hoc Tukey's HSD test |
| S1F | Correlation between Stomatal density and cotyledon area | 14-18 individual cotyledons per accession | 95% CI | Simple linear regression |
| S1G | Correlation between Stomatal Index and Stomatal Density | 8-10 individual cotyledons per accession | 95% CI | Simple linear regression |
| S1H | PCA plot of 9 accessions on 5 environmental variables | Average climatic value for each accession | N/A | Principal Component Analysis |
| S1I | PCA of pairwise genetic distances among all 1135 accessions | 1135 X 1135 data points | N/A | Principal Component Analysis |
| S2A | Simple linear regression of entry ACD vs final SD and final SI, data grouped by accession | bootstrapped, replicate-balanced datasets (n ≈ 14–15 per accession) across six accessions | N/A | Ordinary Least Square (OLS) regression analysis |

| | | | | |
|---|---|---|---|---|
| S2B | $R^2$ values plotted from the simple linear regression, data combined or individually per accession | bootstrapped, replicate-balanced datasets (n ≈ 14–15 per accession) across six accessions | N/A | Ordinary Least Square (OLS) regression analysis |
| S2C | Simple linear regression of entry ACD Vs final SD/SI using the genotype-adjusted means (BLUEs) of each accession | bootstrapped, replicate-balanced datasets (n ≈ 14–15 per accession) across six accessions | N/A | Ordinary Least Square (OLS) regression analysis |
| S2D | Residual plot for entry ACD Vs Final SD, data combined all accessions | bootstrapped, replicate-balanced datasets (n ≈ 14–15 per accession) across six accessions | N/A | Ordinary Least Square (OLS) regression analysis |
| S2E | Residual plot for entry ACD Vs Final SI, data combined all accessions | bootstrapped, replicate-balanced datasets (n ≈ 14–15 per accession) across six accessions | N/A | Ordinary Least Square (OLS) regression analysis |
| S3A | Loading plot of the PCA plot | bootstrapped, replicate-balanced datasets (n ≈ 14–15 per accession) across six accessions | N/A | N/A |
| S3B | Pairwise correlation among stomatal development variables for group1 accessions | bootstrapped, replicate-balanced datasets (n ≈ 14–15 per accession) across three group-I accessions | N/A | Spearman's rank correlation |
| S3C | Pairwise correlation among stomatal development variables for group2 accessions | bootstrapped, replicate-balanced datasets (n ≈ 14–15 per accession) across three group-II accessions | N/A | Spearman's rank correlation |
| S3D | Variable Importance in Projection (VIP) scores for all the accessions together | bootstrapped, replicate-balanced datasets (n ≈ 14–15 per accession) across six accessions | N/A | Partial least Square (PLS) regression model |
| S3E | Variable Importance in Projection (VIP) scores for group1 accessions | bootstrapped, replicate-balanced datasets (n ≈ 14–15 per accession) across three group-I accessions | N/A | Partial least Square (PLS) regression model |
| S3F | Variable Importance in Projection (VIP) scores for group2 accessions | bootstrapped, replicate-balanced datasets (n ≈ 14–15 per accession) across three group-II accessions | N/A | Partial least Square (PLS) regression model |
